# Normal spindle positioning in the absence of EBPs and dynein plus-end tracking in *C. elegans*

**DOI:** 10.1101/118935

**Authors:** Ruben Schmidt, Anna Akhmanova, Sander van den Heuvel

## Abstract

The position of the mitotic spindle is tightly controlled in animal cells, as it determines the plane and orientation of cell division. Interactions between cytoplasmic dynein at the cortex and astral microtubules generate pulling forces that position the spindle. In yeast, dynein is actively delivered to the cortex through microtubule plus-end tracking complexes. In animal cells, an evolutionarily conserved Gα-GPR-1/2^Pins/LGN^–LIN-5^NuMA^ cortical complex interacts with dynein and is required to generate pulling forces, but the mechanism of dynein recruitment to the cortex is unclear. Using CRISPR/Cas9-assisted recombineering, we fluorescently labeled endogenous DHC-1 dynein in *C. elegans.* We observed strong dynein plus-end tracking, which depended on the end-binding protein EBP-2. Complete removal of the EBP family abolished dynein plus-end tracking but not LIN-5-dependent cortical localization. The *ebp-1/2/3* deletion mutant, which was viable and fertile, showed increased cortical microtubule retention; however, pulling forces and spindle positioning were normal. These data indicate that dynein recruited from the cytoplasm creates robust pulling forces.

---

The mitotic spindle determines the plane of cell cleavage through its position and interactions with the cell cortex (reviewed in refs. ^1,2^). The position of the spindle thereby controls the relative size and location of daughter cells formed during cell division. In addition, dependent on the position of the mitotic spindle, polarized cells may divide either symmetrically or asymmetrically. Asymmetric cell divisions generate cell diversity and allow long-term retention of somatic stem cells during the development of organisms and maintenance of tissues. Thus, accurate positioning of the spindle is critical for a wide range of processes that include the formation of gametes, integrity of tissues, creation of different cell types, and coordination of stem cell proliferation and differentiation.

The *C. elegans* early embryo provides an important *in vivo* model for studies of regulated spindle positioning. The one-cell embryo divides asymmetrically based on an anterior-posterior (A-P) polarity axis established after fertilization^3^. Classical spindle severing experiments revealed that pulling forces acting from the cell cortex on astral microtubules (MTs) position the spindle, and that these forces are higher in the posterior than in the anterior^4^. This asymmetry in forces leads to displacement of the spindle towards the posterior, and allows cell cleavage to create two blastomeres of unequal size and developmental fate^5^.

Genetic screens and biochemical experiments have revealed a variety of factors important for the generation and distribution of mitotic pulling forces. Among these are the evolutionarily conserved proteins Gα, GPR-1/2^PINS/LGN^ and LIN-5^Mud/NuMA6–10^. These proteins form a complex known as the force generator complex (FGC), which is regulated downstream of A-P polarity, thus bridging cell polarity and mitotic force generation. Cortical pulling forces also depend on cytoplasmic dynein^11^. This large multi-subunit protein complex is the major minus-end directed motor in the cell and is essential for a large variety of cellular processes, including bipolar spindle assembly, mitotic checkpoint regulation, centrosome positioning, and intracellular transport (reviewed in ref. ^12^). In the current model, the FGC recruits dynein to the cell cortex either directly or indirectly via an extended N-terminal region of LIN-5^13^. Dynein subsequently associates with astral MTs in an end-on configuration, inducing pulling forces via both MT depolymerization and minus-end directed movement. Both *in vivo* and in *vitro* studies support this model^11^,^14–16^.

It remains unclear, however, how dynein is recruited to the cortex and how the dynamic instability of MTs is coupled to force generation and mitotic spindle positioning^17^. MT growth and shrinkage are spatiotemporally modulated by a large variety of MT-associated proteins (MAPs), a subset of which concentrates at the growing MT plus-end (reviewed in ref. ^18^). These MT plus-end-tracking proteins (+TIPs) form a highly interconnected and dynamic network, which is regulated in a cell cycle and position-dependent manner to fine-tune MT dynamics^19–22^. Members of the end-binding (EB) protein family are seen as master regulators of the +TIP network, as they bind autonomously to the growing MT end and recruit multiple other +TIPs^19,23–25^ (reviewed in ref. ^18^).

Interestingly, dynein is known to behave as a +TIP in a variety of cellular contexts^26–29.^ Dynein plus-end tracking in mammals is classically regarded as a ‘search-and-capture’ mechanism, by which the complex finds its cargo molecules via the MT plus-end before initiating minus-end directed transport ^30,31^. In the budding yeast *S. cerevisiae,* dynein plus-end tracking appears coupled to off loading at the cortex and association of dynein with its cortical anchor Num1 ^32–34^. Disruption of dynein plus-end recruitment is thus associated with spindle positioning defects in yeast. Interactions between EB1, CLIP-170, and the dynactin protein p150Glued also recruit the human dynein complex to the MT plus-end^35–37^. However, whether dynein plus-end tracking is generally involved in spindle positioning remains to be determined.

In this study, we explore the localization of dynein during mitotic pulling force generation in the one-cell *C. elegans* embryo. Using CRISPR/Cas9-mediated genome editing^38–42^, we labeled the endogenous dynein complex and studied its dynamic localization during mitosis using a variety of techniques. We observed two populations of dynein at the cortex: an EB-protein dependent MT plus-end tracking dynein population and a LIN-5-dependent MT independent population. By generating single, double and triple EB protein family gene knock-out mutants, we show that spindle positioning and embryonic development can proceed as normal in the absence of dynein plus-end tracking, and despite alterations of MT dynamics.

## Results

### Visualization of the endogenous dynein complex

In order to visualize the dynamics of the dynein complex in the early *C. elegans* embryo we applied CRISPR/Cas9-mediated genome editing. We decided to target the dynein heavy chain as opposed to an accessory subunit in order to label all possible compositions of the dynein complex. In *C. elegans* the dynein heavy chain is encoded by *dhc-1,* which has been labeled with fluorescent proteins (FPs) at both its N- and C-terminus in different systems^43–45^, resulting in targeting of the DHC-1 tail and motor domain, respectively (Fig. 1A). To allow for functional comparison, we explored both strategies by inserting *mcherry* directly upstream of the *dhc-1* stop codon, and *mcherry* or *egfp* directly upstream of the *dhc-1* start codon. A linker region encoding 4 glycine residues was always inserted between FPs and *dhc-1* to preserve independent protein folding (Fig. 1A).

**Figure 1.**
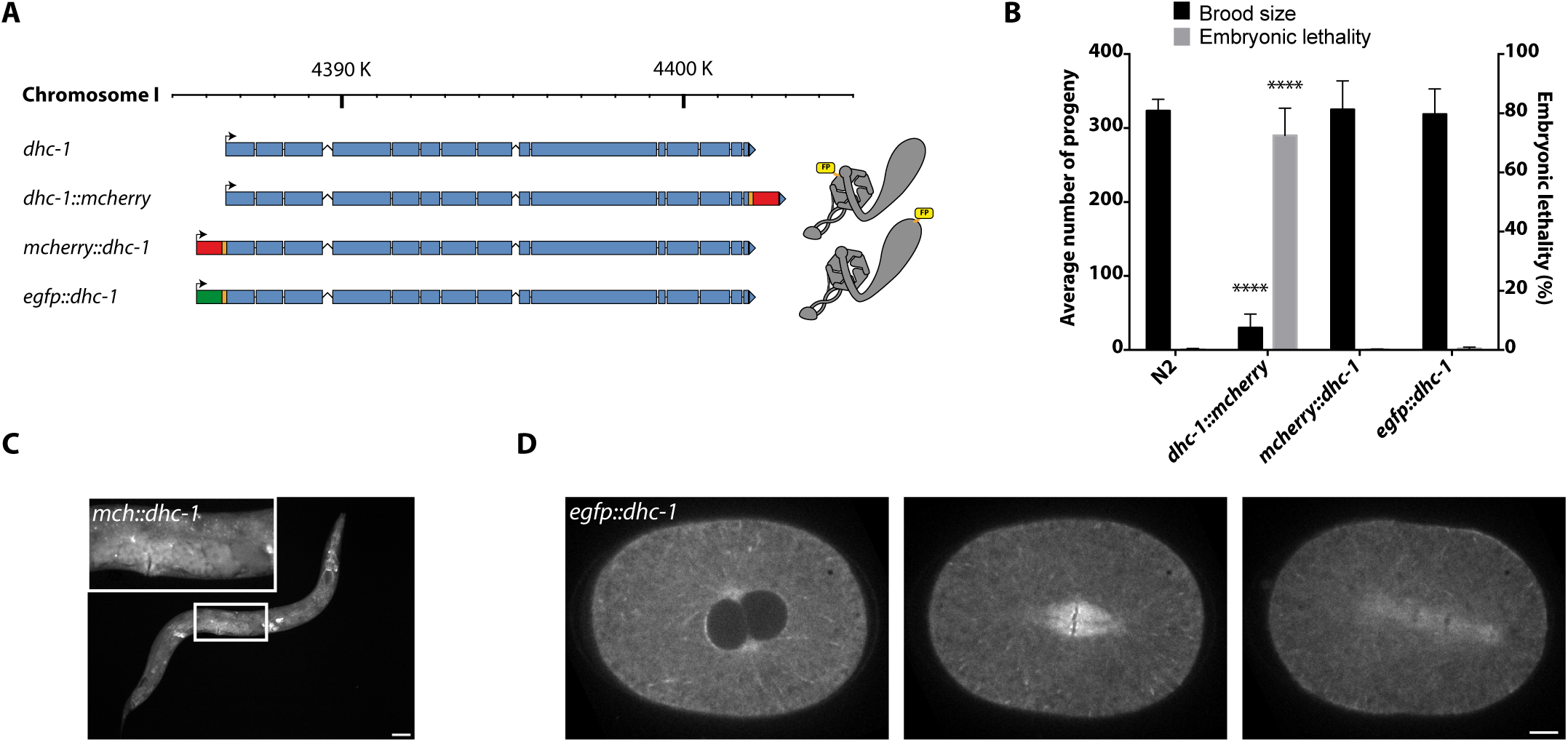
Endogenous tagging of *dhc-1* using CRISPR/Cas9. (A) Schematic representation of *dhc-1* tagging strategies. Blue boxes represent the exons of the *dhc-1* locus on chromosome I. Red and green boxes represent *mcherry* and *egfp* sequences respectively. Orange boxes indicate linkers. Dynein cartoons (right) indicate the location of FPs in DHC-1 for C- (upper) and N-terminal (lower) strategies. (B) Quantification of embryonic lethality (left Y-axis) and brood size (right Y-axis) of *dhc-1* strains. Bars represent average (N=4 replicates) + SD, **** P < 0.0001 compared to wt. No indication means no significant difference to wt. Unpaired Welch Student's *t*-test. (C) Wide-field fluorescence microscopy image of adult animal expressing mCherry::DHC-1. Blow-up indicates expression of mCherry::DHC-1 in the germ-line. Scale bar, 50 μm. (D) Representative spinning disk confocal microscopy images of single eGFP::DHC-1 embryo in prophase (left), metaphase (middle) and anaphase (right). Scale bar, 5 μm. All images were taken with the same exposure time and laser power, anterior to the left.

We obtained multiple homozygous viable knock-in strains using both strategies. However, C-terminal tagging of *dhc-1* resulted in both severe embryonic lethality (72,5% ±9,2 vs. 0,2% ±0,3 in N2), and a strong reduction in brood size (30,0 ±18,4 animals vs. 323,3 ±15,5 in N2) (Fig. 1B). Observation of early embryos by differential interference contrast (DIC) microscopy revealed spindle positioning and cell division defects from the one-cell stage onward. These phenotypes are in accordance with perturbed dynein function^46^, which we confirmed in two independent strains. This led us to conclude that C-terminal tagging of DHC-1 causes a partial loss of DHC-1 function, as a complete loss of function would not be homozygous viable. We did not further study dynein dynamics using these strains.

In contrast, animals with homozygous N-terminally tagged DHC-1 were fully viable (0,1 ± 0,2 in *mcherry::dhc-1* vs. 0,5 ± 0,4 in *egfp::dhc-1* vs. 0,2% ± 0,3 in N2), and showed normal brood size (325,0 ± 38,8 in *mcherry::dhc-1* vs. 318,8 ± 34,2 in *egfp::dhc-1* vs. 323,3 ± 15,5 in N2) (Fig. 1B). Worms appeared healthy during all stages of development, and we observed no obvious abnormalities in early embryonic development for either of these strains by DIC microscopy. From these data we conclude that N-terminal tagging of DHC-1 with either mCherry or eGFP does not perturb dynein function.

Observation of adult worms by wide-field fluorescence microscopy revealed expression of both mCherry::DHC-1 and eGFP::DHC-1 in all somatic tissues of the worm. Notably, DHC-1 is enriched in the germ-line, which is known for the robust silencing of transgenes (Fig. 1C, blow-up). Next we followed early embryos by time-lapse spinning disc confocal laser microscopy (SDCLM). A diffuse cytoplasmic pool of DHC-1 was present in the cytoplasm during all stages of the cell cycle; in addition, DHC-1 was located at the nuclear envelope, centrosomes, kinetochores, kinetochore MTs, central spindle, astral MTs and the cell cortex during mitosis (Fig. 1D). This localization pattern is in accordance with earlier results from immunohistochemistry and transgene overexpression experiments^11,43,46,47.^ In addition, we also noticed comet-like accumulations of dynein radiating from the centrosomes to the cell periphery in a pattern that appeared to follow the astral MT network during metaphase and anaphase.

### Dynein tracks MT plus-ends during mitosis

To further explore this localization pattern, we co-expressed P*pie-1-*driven *gfp:β-tubulin*^48^ with *mcherry::dhc-1* and imaged embryos by simultaneous dual-color time-lapse SDCLM. We observed that dynein comets were MT-associated, most likely concentrating at the growing plus-end (Fig. 2A, arrow, Supplementary Movie 1). Since dynein is a minus-end directed motor protein, we reasoned that cortex-directed dynein comets most likely revealed a plus-end tracking population. To test this possibility, we co-expressed P*pie-1-*driven *ebp-2::gfp* as a germ-line MT plus-end marker^49,50^ with *mcherry::dhc-1*. This revealed strong co-localization of dynein and EBP-2::GFP comets (Fig. 2B, Supplementary Movie 2). Intensity profile analysis of both mCherry::DHC-1 and EBP-2::GFP comets shows strong similarity between the two, indicating that plus-end tracking of dynein is strongly similar to that of EBP-2 (Fig. 2C). Next, we measured the cortex-directed velocity of mCherry::DHC-1 and EBP-2::GFP comets. Measurements indicated that comet velocities are almost identical (0,74 ± 0,11 μm/s for EBP-2::GFP (N=116) and 0,71 ± 0,13 μm/s for mCherry::DHC-1 comets (N=57), p=0,136) (Fig. 2D). These results are in line with earlier quantifications of MT growth speeds during metaphase (0.72 ± 0.02 μm/s^49^). In order to confirm that DHC-1 plus-end tracking is not an artefact due to protein tagging, and to find out whether other components of the dynein complex are present at the MT plus-end, we also studied the localization of dynactin. This protein complex is required for a broad range of dynein functions^12^, one of which is spindle positioning in *C. elegans*^51^. Both GFP::DNC-1^p150glued^ and GFP::DNC-2^p50/dynamitin^ showed clear overlap with mCherry::DHC-1 comets (Supplementary Fig. 1). Thus, we conclude that the dynein comets we observe represent plus-end tracking of the endogenous dynein complex, with high similarity in localization to EB plus-end tracking.

**Figure 2.**
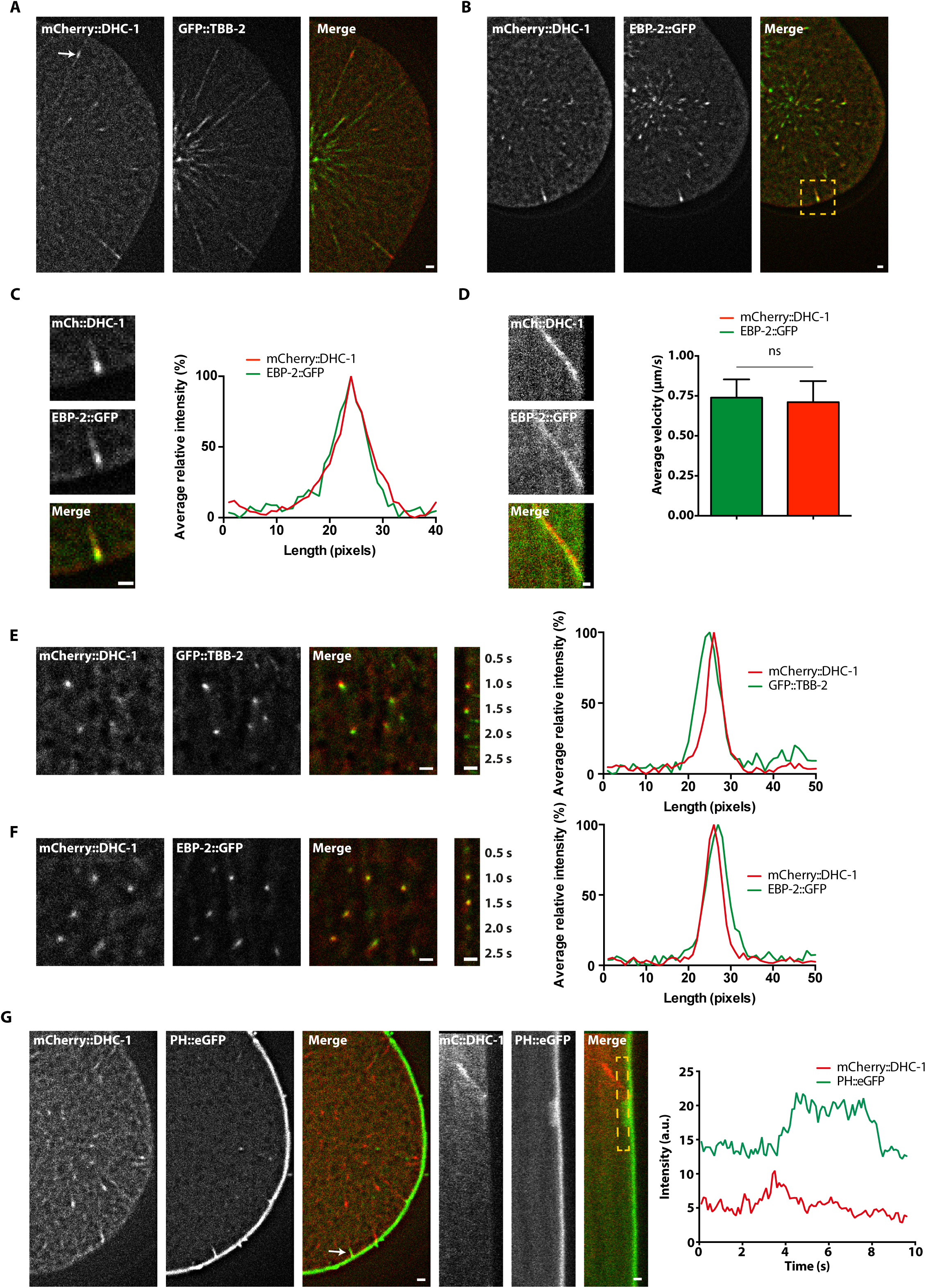
Dynein tracks MT plus-ends during mitosis. (A, B) Representative simultaneous dual-color SDCLM images of one-cell embryos during mitosis. mCherry::DHC-1 (red), GFP::TBB-2 (MTs, green, A), EBP-2::GFP (plus-end, green, B). Arrow indicates dynein plus-end tracking. Yellow box indicates the blow-up depicted in (C). (C) Intensity profile analysis of mCherry::DHC-1 (red) and EBP-2::GFP (green) comets. Length in pixels, intensity in average (N=10 comets) percentage of maximum. (D) Quantification of mCherry::DHC-1 (red, N=57 comets) and EBP-2::GFP (green, N=116 comets) average comet velocity (μm/s) during metaphase. Bars represent average +SD, ns not significant. (E, F) Representative simultaneous dual-color TIRF images of mCherry::DHC-1 and GFP::TBB-2 (tubulin) (E) or EBP-2::GFP (F) localization during early anaphase. Graphs indicate intensity profile of mCherry::DHC-1 (red) and GFP::TBB-2 (E) or EBP-2::GFP (F) decoration of the MT plus-end. Length in pixels, intensity in average (N=10 comets E, F) percentage of maximum. (G) Representative simultaneous dual-color SDCLM images of one-cell embryo expressing mCherry::DHC-1 (red) and PH::eGFP (green, plasma membrane). Panels 4-6 are kymographs of the membrane invagination indicated with the arrow in panel 3. Graph indicates single intensity profile analysis of mCherry::DHC-1 (red) and PH::eGFP as measured along the box delineated in kymograph, representative of N=25 events. Images are averages of 5 consecutive frames taken from 100 ms stream-lapse movies, after background subtraction by a gaussian blur filter. Scale bars, 1 μm.

An important question is how MT plus-end tracking of the dynein complex relates to pulling force generation. Thus, we studied cortical dynein localization in mitosis by dual-color total internal reflection fluorescence (TIRF) microscopy. We observed the simultaneous appearance of dynein comets with end-on MT-cortex contacts at the cortex (Fig. 2E, Supplementary Movie 3). Interestingly, when MTs stopped growing (as judged by loss of EBP-2::GFP signal), we also observed loss of the concentrated mCherry::DHC-1 signal at the cortex (Fig. 2F, Supplementary Movie 4). This indicates that most dynein is lost from the plus-end when the MT switches from a growing to a shrinking state.

To explore the functional relevance of dynein plus-end tracking to spindle positioning, we imaged dynein dynamics during cortical force generation. Single force generation events can be visualized by invaginations of the plasma membrane^52^. To visualize membrane invaginations, we used CRISPR/Cas9-mediated genome editing to generate a strain expressing PH::eGFP driven by *Peft-3* in all tissues including the germ-line. This transgene was integrated in the *CxTi10816* locus on chromosome IV previously used to achieve germ-line expression of single-copy integrated transgenes^53^. Interestingly, dual-color imaging of PH::eGFP and mCherry::DHC-1 reveals strong co-occurrence of dynein comets reaching the cortex and the formation of membrane invaginations in 13/25 events (Fig. 2G, Supplementary Movie 5). However, we could not detect enrichment of mCherry::DHC-1 after the initiation of membrane invaginations. This indicates that a significant part of the dynein population present in a comet at the MT plus-end dissipates upon membrane contact and is most likely not involved in the generation of cortical pulling forces.

### Dynein plus-end tracking and *ebp-1/2/3* are not required for spindle positioning and development

Although disappearing during membrane invagination, it remained unclear whether the MT plus-end tracking population of dynein contributes to cortical pulling force generation. To investigate this possibility, we sought to specifically disrupt dynein plus-end accumulation by removing +TIPs. To date, MT plus-end tracking has not been thoroughly explored in *C. elegans,* though two homologs of the mammalian EBs, EBP-1 and EBP-2, have been shown to exhibit plus-end tracking activity when overexpressed in the embryo^49^. An annotated third homolog, *ebp-3,* appears to have arisen by a duplication of the genomic region containing the *ebp-1* gene. This duplication likely occurred relatively recently, as judged by the almost complete sequence identity between *ebp-1* and *ebp-3*. However, alignment of the predicted protein sequences revealed that EBP-3 lacks the calponin homology (CH) domain, which is required for recognition of the MT plus-end^24,54.^ Thus, we suspect that *ebp-3* is a pseudogene. Because of the high sequence similarity between *ebp-1* and *ebp-3,* we refer to these genes collectively as *ebp-1/3*.

We knocked down expression of *ebp-1/3* and ebp-2 by RNAi, and imaged one-cell *egfp::dhc-1* embryos during mitosis by time-lapse confocal microscopy. While loss of *ebp-1/3* expression did not appear to affect eGFP::DHC-1 plus-end tracking, loss of *ebp-2* expression abolished the appearance of dynein comets during mitosis (Fig. 3A). Interestingly, this did not result in obvious spindle positioning or cell division defects upon initial inspection.

**Figure 3.**
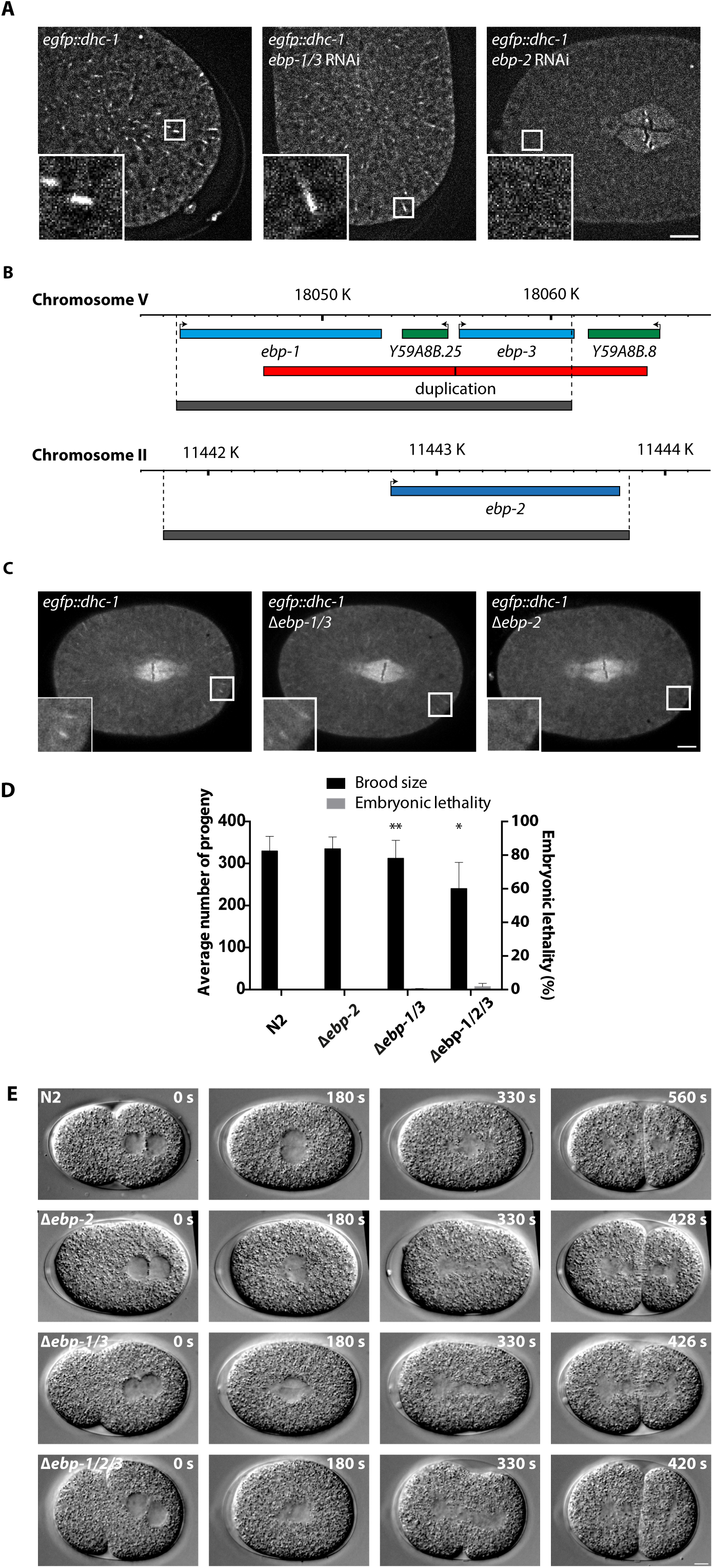
Analysis of dynein plus-end tracking and development in Δ*ebp* knock-outs. (A) Representative SDCLM images of one-cell embryos expressing eGFP::DHC-1 during metaphase, either untreated (wt, left) or following RNAi of indicated *ebp* gene(s) (middle, right). Blow-ups indicate the presence or absence of MT plus-end tracking, which was confirmed in N=5 embryos per condition. Images are averages of 5 consecutive frames taken from 100 ms stream-lapse movies, after background subtraction by a gaussian blur filter. (B) Schematic representation of *ebp-1/3* and *ebp-2* endogenous knock-out strategies. Blue and green boxes represent relevant genes mentioned in the text. Red boxes indicate the genetic duplication on chromosome V mentioned in the text. Grey bars represent the Δ*ebp-1/3* and Δ*ebp-2* deletions. (C) Representative SDCLM images of one-cell embryos expressing eGFP::DHC-1 during mitosis, in a wt (left), Δ*ebp-1/3* (middle) or Δ*ebp-2* (right) genetic background. Blow-ups indicate the presence or absence of MT plus-end tracking, which was confirmed in N=5 embryos per condition. Images are selected frames from 1000 ms exposure time-lapse movies, processed by background correction. (D) Quantification of embryonic lethality and brood size of Δ*ebp* mutants. Bars represent average (N=4 replicates) + SD, * P < 0.05, ** P < 0.01 compared to wt. Unpaired Welch Student's *t*-test. (E) Representative single images taken from time-lapse DIC movies of N2, Δ*ebp-2*, Δ*ebp-1/3* and Δ*ebp-1/2/3* one-cell embryos, aligned in time with PNM (first), centration (second), PSD (third) and cytokinesis completion (fourth column) in wild-type (N2). Times relative to pronuclear meeting are indicated in seconds for every frame. Scale bars, 5 μm.

To further explore the role of the EBPs and dynein plus-end tracking during mitosis, and to circumvent the possibility of incomplete RNAi knockdown, we generated knock-out alleles for both *ebp-2* and *ebp-1/3* (Fig. 3B). Making use of CRISPR/Cas9, the neighboring *ebp-1* and *ebp-3* genes were removed together, by excising a ~17 kb region (referred to as Δ*ebp-1/3*). The *ebp-2* knock-out allele lacks both the coding region as well as ~1 kb upstream of the start codon (referred to as Δ*ebp-2).* Both the Δ*ebp-1/3* and Δ*ebp-2* strains could be stably maintained as homozygotes, indicating that neither *ebp-1/3* nor *ebp-2* are strictly required for normal development and reproduction.

When crossed with *egfp::dhc-1,* we observed effects on dynein localization identical to those of the RNAi experiments. Dynein still localized to the MT plus-end in Δ*ebp-1/3; egfp::dhc-1* embryos, while this localization was completely absent in the Δ*ebp-2; egfp::dhc-1* early embryos (Fig. 3C, Supplementary Movie 6). At the same time, dynein localization to the nuclear envelope, spindle midzone and poles did not appear to be affected in either case, which indicates that dynein depends on EBP-2 specifically for recruitment to astral MT plus-ends (Supplementary Fig. 2).

While generally showing no defects in development, Δ*ebp-1/3* caused a slight but statistically significant reduction in brood size (312,50 ± 42,53 in Δ*ebp-1/3* vs. 329,75 ± 35,11 in N2), while Δ*ebp-2* did not (335,25 ± 28,04 in Δ*ebp-2* vs. 329,75 ± 35,11 in N2). In addition, Δ*ebp-1/3,* but not Δ*ebp-2,* displayed a significant but biologically probably irrelevant increase in embryonic lethality (0,47% ± 0,16 in Δ*ebp-1/3* and 0,00% ± 0,00 in Δ*ebp-2* vs. 0,07% ± 0,14 in N2) (Fig. 3D). In order to assess whether development and reproduction could proceed in the absence of all three EBPs, we crossed Δ*ebp-1/3* with Δ*ebp-2* to generate the triple knock-out mutant Δ*ebp-1/2/3.* Interestingly, this combination was viable and could be maintained as triple homozygous animals. The triple mutant showed a reduction in brood size (240,38 ± 62,45 vs. 329,75 ± 35,11 in N2), which is stronger than the observed reduction in Δ*ebp-1/3* animals and indicates some redundancy amongst *ebp-1/3* and *ebp-2.* Embryonic lethality remained low (1,79% ± 1,96 vs. 0,07% ± 0,14 in N2), which is remarkable given the complete absence of EBPs and expected profound disruption of the +TIP network. While Δ*ebp-1/2/3* worms generally show no developmental defects, we did observe a partly penetrant pleiotropic phenotype amongst adult worms. This included low frequencies of dumpy, sterile, and/or uncoordinated animals, as well as lethal larvae that exploded through the vulva. In addition, some triple mutant worms appeared to form small bulges in the skin. Because of the low penetrance and unpredictable nature, we did not further examine these abnormalities.

In order to assess whether spindle positioning was affected in the Δ*ebp* mutants, we followed early embryos by DIC time-lapse microscopy and characterized key mitotic events. Importantly, asymmetric positioning of the spindle and subsequent asymmetric division of the one-cell embryo was not affected in Δ*ebp-1/3,* Δ*ebp-2* or Δ*ebp-1/2/3* embryos (Supplementary Fig. 3A). Δ*ebp-1/2/3* embryos did exhibit a slight change in geometry, as they attained a rounder shape compared to wild-type (wt) embryos (Supplementary Fig. 3B). The position of pronuclear meeting along the A-P axis of the embryo did not change significantly (Supplementary Fig. 3C), while centration of the nucleocentrosomal complex occurred slightly more posterior in Δ*ebp-1/3,* Δ*ebp-2* and Δ*ebp-1/2/3* embryos. The angle at which the metaphase spindle is set up did not change significantly (Supplementary Fig. 3E), while elongation of the anaphase spindle was slightly increased only in the Δ*ebp-1/2/3* mutant (Supplementary Fig. 3F). Regardless of these very small variations in mitotic events, they did not appear to have any significant impact on the outcome of mitosis. However, all Δ*ebp* strains did exhibit an accelerated progression through mitosis compared to wt embryos (Fig. 3E and Supplementary Fig. 4A, Supplementary Movie 7). This effect occurred in Δ*ebp-1/3* and Δ*ebp-2* mutants, but was most dramatic in Δ*ebp-1/2/3.* Both the time to progress from pronuclear meeting to nuclear envelope breakdown (NEBD) (Supplementary Fig. 4B), as well as from the start of chromosome segregation until completion of furrow ingression was significantly decreased (Supplementary Fig. 4D). We do not know the mechanism behind the shortened M phase. However, there was no significant reduction in the time between NEBD and anaphase onset (Supplementary Fig. 4C), which indicates that the observed effect does not result from bypassing or premature satisfaction of the spindle assembly checkpoint.

**Figure 4.**
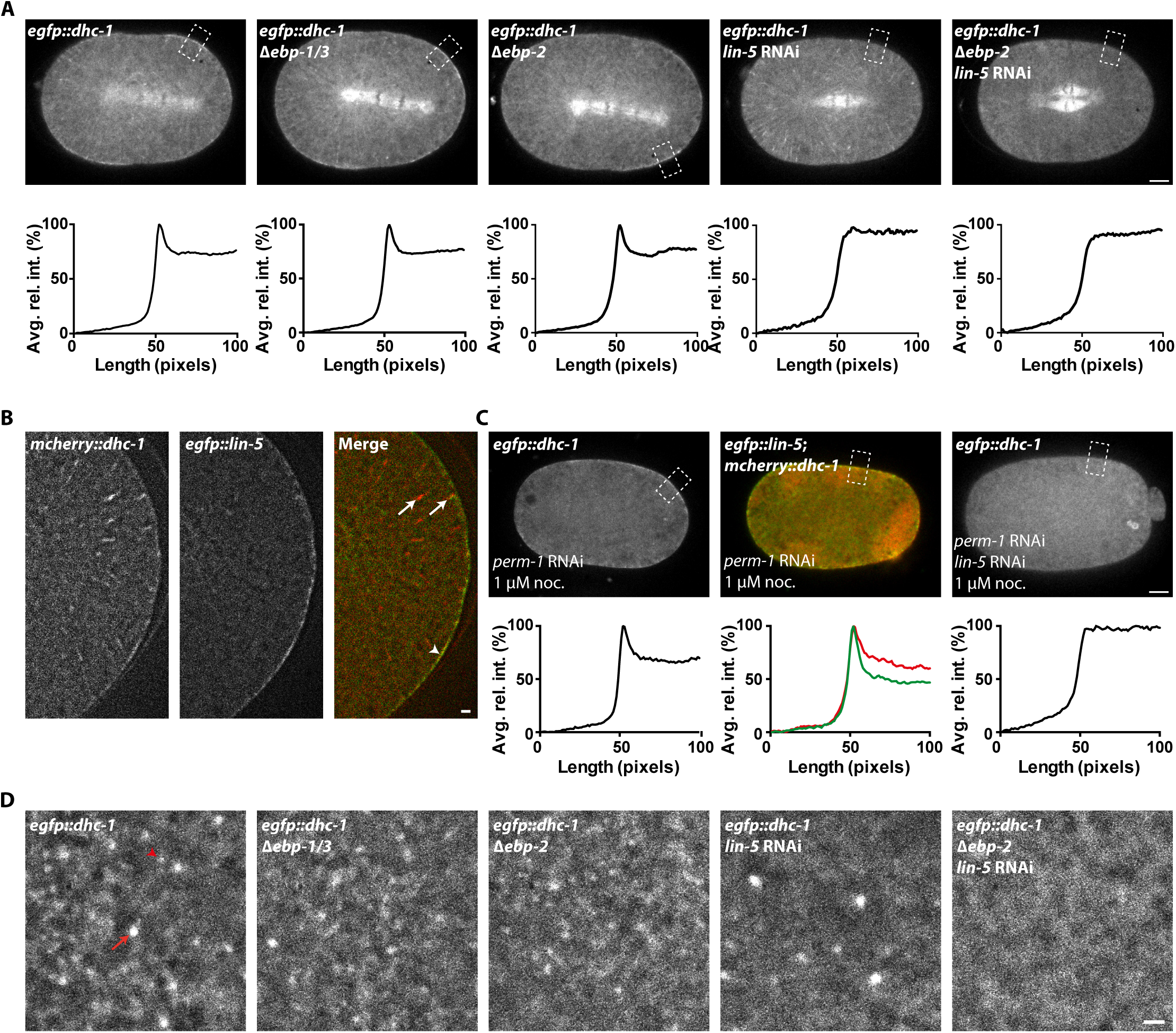
Cortical localization of dynein depends on LIN-5, but not on EBP-2- mediated plus-end tracking. (A) Representative SDCLM images of eGFP::DHC-1 localization during anaphase, under different RNAi conditions and in various genetic backgrounds as indicated for each panel. Graphs indicate intensity profiles of eGFP::DHC-1 as measured in boxes indicated in each superimposed panel. Length in pixels, intensity relative to cytoplasmic values in average (N=24 measurements, 4 embryos per condition) percentage of maximum. Images are selected frames from 1000 ms exposure time-lapse movies, processed with background correction. Scale bar, 5 μm. (B) Representative simultaneous dual-color SDCLM images of mCherry::DHC-1 (red) and eGFP::LIN-5 (green) localization during mitosis. Arrow indicates dynein plus-end tracking, arrowhead indicates co-localization of dynein and LIN-5 at the cortex, as confirmed in N=5 embryos. Images are averages of 10 consecutive frames taken from 100 ms stream-lapse movie, after background subtraction by a gaussian blur filter. Scale bar, 1 μm. (C) Representative SDCLM images of either eGFP::DHC-1 or mCherry::DHC-1; eGFP::LIN-5, in embryos treated with *perm-1* or *perm-1* + *lin-5* RNAi, and with 1 μm nocodazole. Graphs indicate intensity profiles of eGFP::DHC-1 or mCherry::DHC-1 (red) and eGFP::LIN-5 (green) as measured in boxes delineated in each superimposed panel. Length in pixels, intensity relative to cytoplasmic values in average (N=10 measurements, 10 embryos per condition) percentage of maximum. (D) Representative TIRF images of cortical eGFP::DHC-1 during early anaphase, under different RNAi conditions and in various genetic backgrounds as indicated for each panel. Arrow indicates large dot (plus-end dynein), arrowhead indicates small dot (cortical dynein), as discussed in the text and observed for N=6 embryos per condition. Images are averages of 10 consecutive frames taken from 50 ms stream-lapse movie, after background subtraction by a gaussian blur filter. Scale bar, 1 μm. Scale bar, 5 μm.

### LIN-5 recruits dynein to the cell cortex independently of its ability to track the MT plus-end

If MT plus-end tracking of the dynein complex is not required for spindle positioning, then how does dynein reach the cortex? To find out, we assessed the cortical localization of dynein in both wt and Δ*ebp* mutants. Previous studies have attempted to visualize cortical dynein localization by immunohistochemistry^11,13.^ While cortical dynein could be observed in two-cell embryos, immunostaining of one-cell embryos did not show clear cortical localization. Time-lapse SDCLM of *egfp::dhc-1* one-cell embryos revealed broad and transient regions of DHC-1 enrichment at the cortex during anaphase. These were most pronounced during rocking of the spindle. Interestingly, this cortical enrichment of dynein was not perturbed in Δ*ebp-1/3* or Δ*ebp-2* mutant embryos (Fig. 4A). Thus, we conclude that EBP-1/3, and EBP-2-dependent plus-end tracking are not required for the cortical localization of dynein. By contrast, knock-down of *lin-5* perturbed cortical dynein localization but did not affect its localization to MT plus-ends (Fig. 4A). Simultaneous dual-color imaging of embryos expressing endogenously-tagged *egfp::lin-5* and *mcherry::dhc-1* showed that dynein comets do not co-localize with LIN-5, whereas cortical dynein and LIN-5 co-localize at the cortex (Fig. 4B). Moreover, we previously observed cortical dynein localization in the absence of MTs, following treatment of *perm-1(RNAi)* embryos with 1 μM nocodazole (Portegijs *et al.,* accepted manuscript). In the absence of astral MTs, as confirmed by tubulin and EBP-2 markers, dynein still localized to the cortex, and this localization strongly overlapped with and depended on LIN-5 (Fig. 4C). In addition, we visualized cortical dynein in the early embryo by time-lapse TIRF microscopy. Notably, this revealed two populations of dynein that could be genetically separated at the cortex: a population of bright large spots that represent dynein comets at the cell cortex (Fig. 4D, arrow), and a fraction of smaller, less dynamic spots that represent a LIN-5 dependent population (Fig. 4D, arrowhead). Most notably, the combination of Δ*ebp-2* with knock-down of *lin-5* resulted in a complete loss of cortical dynein (Fig. 4D, Supplementary Movie 8). Thus, two independent populations of dynein appear present at the cortex: an EBP-2-dependent plus-end tracking and a LIN-5-dependent cortical population. LIN-5 can most likely recruit dynein directly from the cytoplasm to the cortex, which would explain why plus-end tracking of dynein is not required for its cortical localization.

### Cortical pulling forces remain normal in the absence of end-binding proteins and dynein plus-end tracking

To assess the effect of the absence of EBPs and dynein plus-end tracking on anaphase pulling forces, we imaged embryos by DIC time-lapse microscopy during mitosis. Subsequently, we tracked anterior and posterior pole movements during anaphase and calculated the average maximum amplitude, which is a read-out for force generation during anaphase^55^. Interestingly, rocking of both the anterior and posterior centrosome was generally not affected by the removal of EBPs. We did find a slight but significant increase in the maximum amplitude for the posterior pole in the Δ*ebp-1/3* background, but this change was not present in the Δ*ebp-1/2/3* background (Fig. 5A).

**Figure 5.**
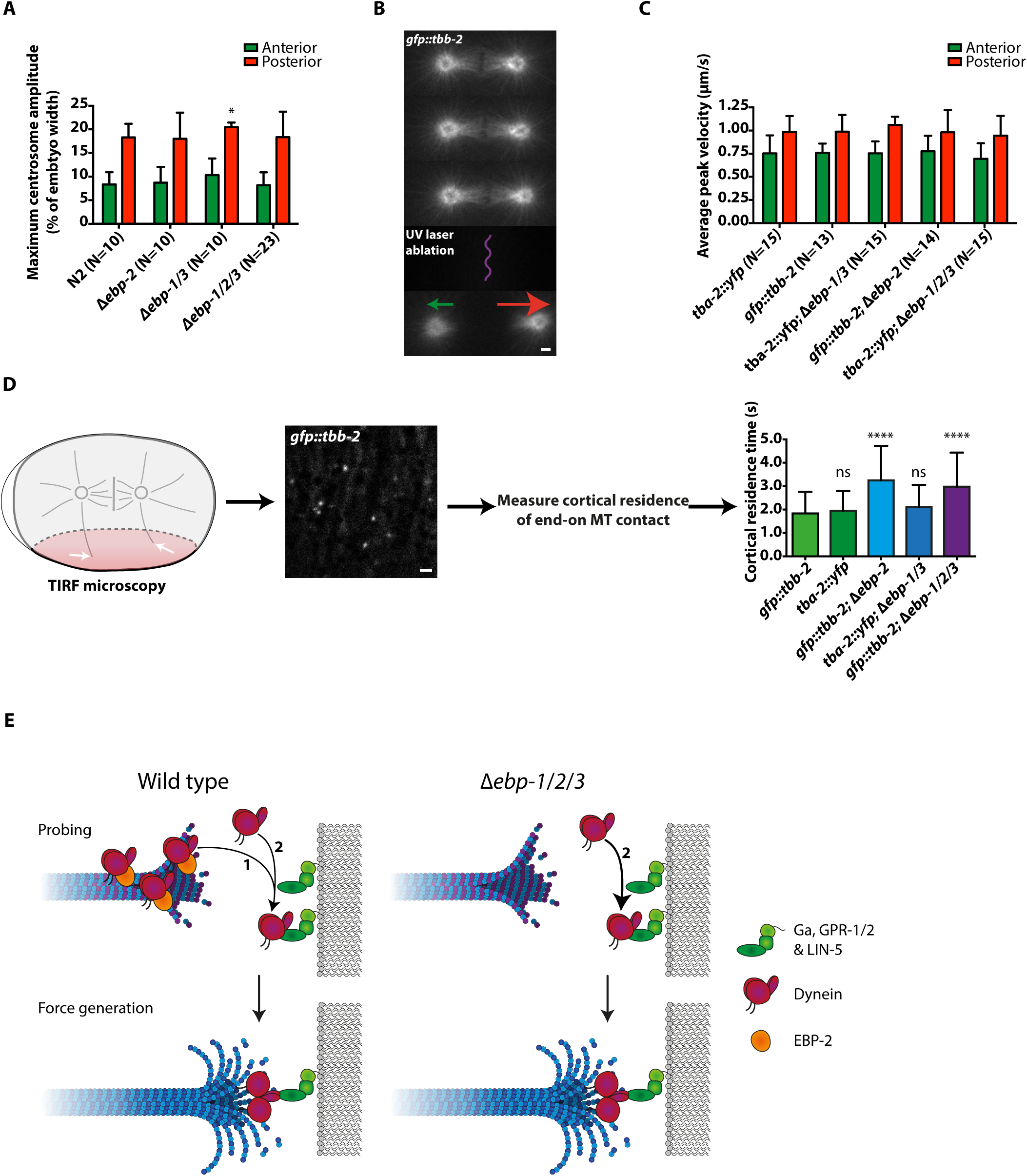
Loss of EBP-1/2/3 affects MT dynamics but does not affect mitotic pulling force generation. (A) Quantification of centrosome rocking during anaphase in wt and Δ*ebp* embryos. The max. amplitude is presented as a % of embryo height for both the anterior (green) and posterior (red) pole. Bars represent average + SD, * P < 0.05 compared to wt. Unpaired Welch Student's *t*-test. N-values are indicated below bar graphs. (B) Example of spindle bisection using an ultraviolet laser (curved purple line) in an embryo expressing GFP::TBB-2 (MTs). Five consecutive frames are shown, each the average of 2 consecutive frames from a 500 ms exposure stream-lapse movie. Arrows indicate the direction and arbitrary relative speed of displacement upon spindle bisection. Anterior to the left. Scale bar, 1 μm. (C) Quantification of average peak velocity (μm/s) upon spindle bisection for anterior (green) and posterior (red) centrosomes. Bars represent average + SD, no indication means no significant difference from wt. Unpaired Welch Student's *t*-test. N-values are indicated below bar graphs. (D) Quantification of cortical residence time of end-on MT contacts as visualized by TIRF microscopy of one-cell embryos expressing either GFP::TBB-2 or TBA-2::YFP during early anaphase (schematically represented on the left), in different Δ*ebp* mutant backgrounds. Representative image shown is an average of 5 consecutive frames from a 250 ms stream-lapse movie, after background subtraction by a gaussian blur filter. Scale bar, 1 μm. Average cortical residence time in seconds. Bars represent average + SD, **** P < 0.0001 compared to wt, ns not significant. Unpaired Welch Student's *t*-test. N=100 tracks for each condition. (E) Model of force generation in wt (left) and Δ*ebp-1/2/3* (right) as explained in the text.

To quantify the forces generated during anaphase more directly, we performed spindle severing assays with a focused UV laser beam ^4^. Upon severing of the central spindle during anaphase onset, the centrosomes move away from each other with a velocity proportional to the pulling forces acting on their astral ΜTs ^4^ (Fig. 5B, Supplementary Movie 9). Remarkably, we did not find any significant differences in average peak velocity between any of the Δ*ebp* mutants and wild-type embryos for either pole (Fig. 5C). Thus, cortical pulling force generation is robust and remains largely unaltered in the absence of EB proteins. We conclude that dynein plus-end tracking is not required for cortical pulling force generation in the one-cell *C. elegans* embryo.

The EB proteins not only determine dynein plus-end tracking but are also thought to affect several aspects of MT dynamics. The direct effects of EBs on MT dynamics are still unclear, because *in vitro* and *in vivo* studies have yielded contradictory results^25,56–58^. The prevailing view is that EBs increase the growth rate as well as the catastrophe frequency of MTs *in vitro,* but have little effect on the growth rate and suppress catastrophes in cells. However, the effect of complete loss of EB family members on MT dynamics has not been reported in an established *in vivo* system such as *C. elegans.* A hurdle for studying MT dynamics in the one-cell embryo is the extremely dense MT network, with each centrosome of the spindle concurrently nucleating about 300 MTs^50^. This complicates the observation of single MTs by GFP-tubulin labeling, and we could not use +TIP markers as an alternative in this case. Therefore, we imaged the dynamics of labeled tubulin at the cortex with the use of TIRF microscopy. By quantification of the end-on MT-cortex contacts during early anaphase, we found a significant increase in residence time in both Δ*ebp-2* and Δ*ebp-1/2/3,* but not for Δ*ebp-1/3,* mutants compared to wild type (Fig. 5D, Supplementary Movie 10). This indicates that loss of EBP-2 either reduces MT growth rate, catastrophe frequency or both, thereby possibly allowing prolonged contact with the cell cortex. Interestingly, we also observed a reduced ΜT density of the spindle midzone in Δ*ebp-2* and Δ*ebp-1/2/3* embryos (Supplementary Fig. 5). This led to full or partial bisection of spindles, reminiscent of *spd-1^PRC1^* knock-down, which diminishes the mechanical strength required for the midzone to counteract the forces acting on the centrosomes^59^. We attribute this effect to a potential reduction in MT polymerization. Collectively, these results indicate that the +TIP network composition, dynein plus-end tracking, and MT dynamics are altered by the loss of the EBPs. However, the net pulling forces are not affected, possibly due to compensation by altered MT dynamics.

## Discussion

In this study, we investigated the recruitment of dynein in the generation of pulling forces that position the mitotic spindle and determine the plane of cell division. We took advantage of the well-established *C. elegans* one-cell embryo as an *in vivo* model for spindle positioning, by combining targeted genome editing with high spatial and temporal resolution microscopy, spindle severing assays, RNAi, and drug treatment. Using CRISPR/Cas9-assisted recombineering, we tagged the endogenous dynein complex and created knockout alleles for all three genes encoding plus-end binding proteins of the EB1 family. The ability to study endogenously expressed proteins allowed for reliable phenotypic characterizations and observation of protein dynamics without overexpression.

Our first observation was that C-terminal tagging of the *dhc-1* dynein heavy chain causes partial loss of function, as opposed to N-terminal tagging. It is currently unclear whether this translates to dynein in other organisms. N-terminally and C-terminally tagged yeast Dyn1 were reported to be functional^60^. However, Dyn1 lacks the C-terminal regulatory extension present in the cytoplasmic dynein heavy chain of other organisms^61^ including *C. elegans.* A BAC transgene with a C-terminally GFP-tagged mouse dynein heavy chain is commonly used in mammalian studies^62^, normally in the presence of the endogenous DHC protein and in cells in culture, which may depend less critically on dynein function. Tagging the N-terminal DHC tail region instead of the C-terminal motor domain might be the best option for future *in vivo* studies.

The next interesting observation was the concentration of the dynein complex at the growing MT plus-ends in mitosis. This was previously described for other systems, but to our knowledge not in *C. elegans.* The detection of dynein plus-end tracking allowed us to quantify whether this transient dynein localization forms part of the mechanism for cortical pulling force generation. Surprisingly, pulling forces and spindle positioning remained unaltered in the absence of dynein plus-end tracking and removal of the entire EB protein family. These observations are in clear contrast with data from budding yeast, which indicate that plus-end tracking provides an active system to deliver dynein to the daughter cell cortex.

*In vivo* and *in vitro* experiments with *S. cerevisiae* Dyn1 have implicated the EB1-related protein Bim1^EB1^, together with Bik1^CLIP170^, Pac1^LIS1^, and Kip2^Kinesin^ in dynein plus-end recruitment^63,64.^ Yeast dynein is thought to be primed at the MT plus-end for cortical anchorage by relieving its auto-inhibition^32,33,65.^ Offloading from the MT plus-end to the cortex is dynactin-dependent, and allows cortical anchorage by Num1 and correct spindle positioning^32,34.^ This two-step mechanism was suggested to locally recruit dynein while at the same time minimizing the need for modulation of motor activity.

In yeast, an estimated 30% of plus-end dynein appears delivered via kinesin-mediated transport, while 70% is recruited from the cytoplasm^34^. This, together with the finding that Bik1^CLIP170^ can recruit dynein to the MT plus-end in absence of Bim1^EB1 66^, likely explains why depletion of Bim1^EB1^ from yeast cells does not completely abolish dynein plus-end tracking^63^. Isolated Dyn1 shows constitutive minus-end-directed processive movement upon dimerization *in vitro^60^.*

This indicates why kinesin plus-end directed transport is important for keeping Dyn1 at the plus-end^64^. While we did not explore the role of kinesin motors in the context of dynein plus-end tracking, the effect of EBP-2 depletion suggests that kinesin-mediated transport of dynein does not play a major role in *C. elegans.* Possibly, *C. elegans* dynein regulation is more related to the human than to the yeast dynein complex.

*In vitro* reconstitution studies have shown that the human dynein complex is recruited to MT plus-ends by the large dynactin subunit p150Glued, which binds to growing MT ends through EB1 and CLIP-170^37^. As a major difference, yeast dynein dimers are active on their own, while mammalian cytoplasmic dynein is in an inactive conformation and requires adaptor-mediated dynactin binding for processive movement^67, 68.^ If both human and *C. elegans* dynein are kept inactive at the plus-end, then the ‘tug-of-war’ with a potential kinesin should not be needed. Kinesin and LIS1 dependent dynein transport has been observed, but its role may be specific for certain cell types such as neurons^69^. However, a recent study indicates that LIS-1 is required for dynein plus-end tracking in *C. elegans* (Rodriguez Garcia *et al.,* submitted). Interestingly, subcellular location-dependent adaptor proteins bridge dynein and dynactin to activate dynein processivity, including the coiled-coil proteins Bicaudal D at intracellular membranes, Rab11-FIB3 at Rab11-endosomes and Spindly at the kinetochore. It is attractive to hypothesize that LIN-5/NuMA acts as a dynein-activating adaptor at the cell cortex.

Could MT plus-end tracking of dynein play a role in asymmetric force generation? In *S. cerevisiae,* dynein cortical recruitment is polarized by its ability to track the MT plus-end, which directs dynein transport to the bud cortex^32^. However, our data would suggest that dynein plus-end tracking is not in itself a polarizing mechanism in *C. elegans.* We did not observe any obvious asymmetry in dynein localization to the MT plus-end, and the spindle as well as the distribution of end-on MT contacts were shown to be symmetric in the one-cell embryo^50^. In addition, the presence of cortical dynein and normal spindle positioning in the absence of EBPs indicates that dynein plus-end tracking is not required for asymmetric spindle positioning. This is further supported by data from nocodazole treatment of embryos, which strongly suggests that LIN-5 can recruit dynein directly from the cytoplasm, as opposed to MT-mediated delivery of dynein to the cortex. This is in accordance with observations in nocodazole-treated cultured mammalian cells, which have shown that cortical dynein recruitment does not require MT polymerization. At the same time, the correct distribution of cortical dynein has been reported to depend on a dynamic astral MT network^70^. Thus, in *C. elegans* and mammalian cells, MT-mediated delivery does not appear necessary for dynein localization to the cortex, but dynamic MTs appear generally required for the correct distribution of dynein at the cortex.

Based on our results, we propose that cortical recruitment of dynein in *C. elegans* occurs by a mechanism different from the off-loading in yeast. We did not observe off-loading or minus-end directed transport of dynein when MT plus-ends reach the cortex. Although rapid diffusion appeared to prevent visualization, the strong similarity and dependence in localization supports that dynein follows EBP-2 plus-end association and release. Intriguingly, we found that pulling force generation and spindle positioning are completely normal in the absence of EBPs. This was surprising, as we did observe effects on MT dynamics and +TIP network composition. It is worth noting that a minor dampening of centrosome rocking amplitudes was observed upon perturbation of *ebp-2* in a recent study complementary to ours (Rodriguez Garcia *et al.,* submitted). Our quantifications do not reveal such an effect in Δ*ebp-2* embryos, which might be explained by the use of different methods of quantification, reporter strains and mutant alleles.

How could the embryo compensate for the observed effects on MT dynamics, and what is the relevance of dynein plus-end tracking in the context of spindle positioning? We propose that in a normal situation, dynein plus-end tracking could function as a local enrichment mechanism (Fig. 5E). By concentrating dynein at the plus-end, MTs could efficiently constitute complete FGCs with complexes of Gα-GPR-1/2–LIN-5 waiting at the cortex (Fig. 5E, arrow 1). This local concentration of dynein could also increase the relative amount of complete FGCs present at the cortex. Thus, dynein plus-end tracking could be a back-up mechanism that ensures efficient force generation. In the absence of EBPs and thus dynein plus-end tracking, astral MTs would have to locate cortical complexes that anchor dynein directly from the cytoplasm (Fig. 5E, arrow 2). While this single mechanism was previously hypothesized to be less efficient^71^, we expect that the prolonged cortical residence of MTs might allow for successful probing of the cortex. In addition, considering that the +TIP is a protein-dense network in which a limited amount of binding sites are available at any given time, cortical FGCs may associate with MTs more efficiently due to decreased crowding of MT tips in EB-depleted cells. Taken together, our work illustrates the complexity and robustness of molecular mechanisms controlling an essential cellular process such as spindle positioning.

## Materials and Methods

### *C. elegans* strains

A summary of the strains used in this study is included in Table 1. All strains were maintained at 20 °C as described previously^72^, unless stated otherwise. Worms were grown on plates containing nematode growth medium (NGM) seeded with OP50 *Escherichia coli* bacteria.

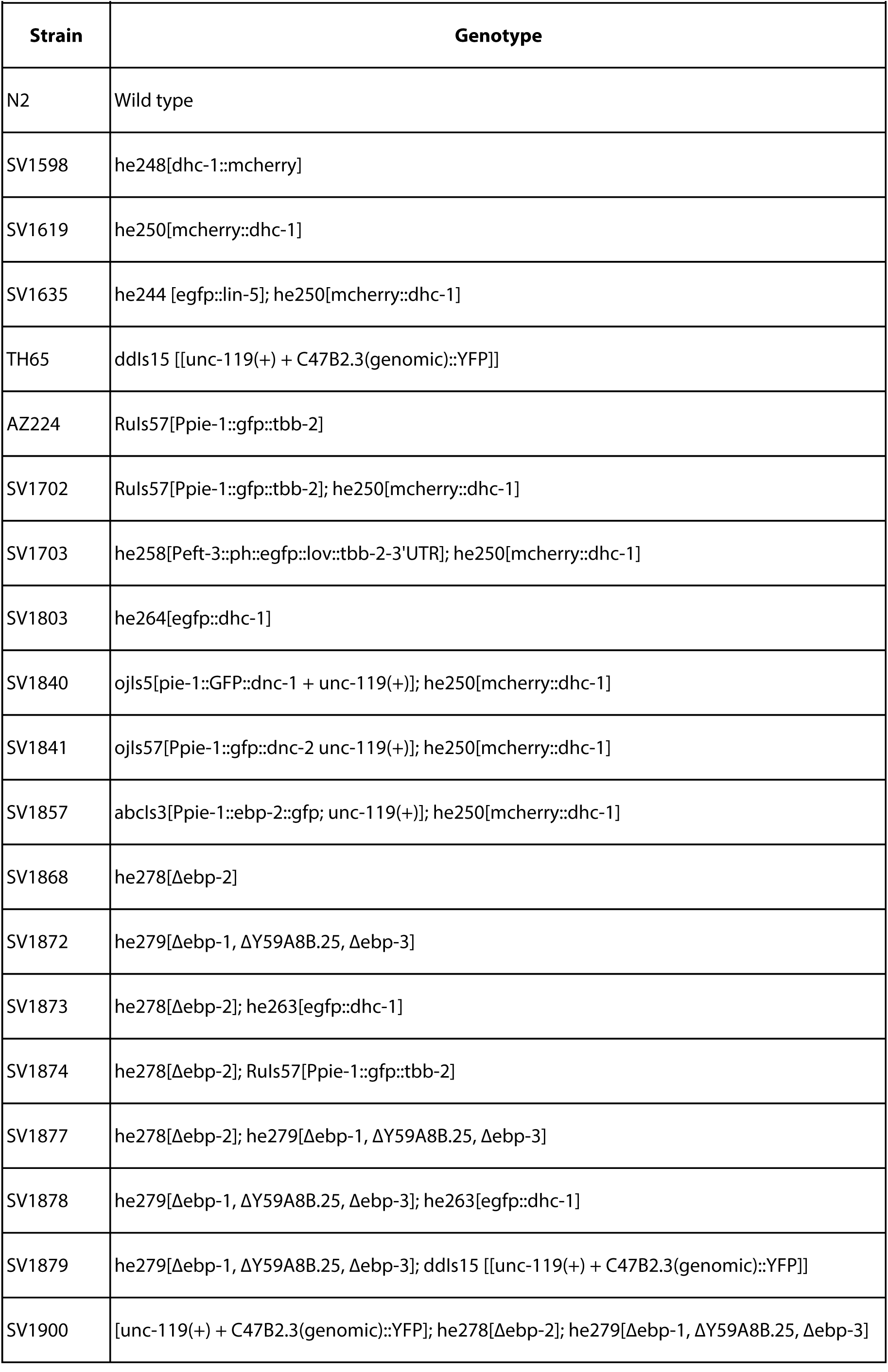

### Generation of CRISPR/Cas9 repair templates and gRNAs

Homology arms of at least 1500 bp flanking the CRISPR/Cas9 cleavage site were generated by PCR amplification from purified *C. elegans* genomic DNA using the KOD polymerase (Novagen). PCR products were inserted into the pBSK backbone by Gibson assembly (New England Biolabs). For the generation of *ph::egfp::lov* and *egfp::dhc-1, egfp* was amplified from pMA-*egfp, ph* from *Pwrt-2::gfp::ph*^73^ and *lov* from *gfp::LOVpep::unc-54UTR*^74^. For *mcherry::dhc-1* and *dhc-1::mcherry,* codon-optimized *mcherry* was amplified from TH0563-PAZ-*mCherry* (a kind gift from A. Hyman). Primers containing overlaps between PCR fragments, linker sequences and mutated gRNA sites were synthesized (Integrated DNA technologies). For the generation of gRNA vectors, oligonucleotides were annealed and inserted into pJJR50 using T4 ligation (New England Biolabs). Vectors were used to transform and purified from DH5alpha competent cells (Qiagen).

### CRISPR/Cas9-mediated genome editing

Injection of adult *C. elegans* worms in the germ-line was carried out using an inverted microscope micro-injection setup. Injection mixes contained a combination of 30-50 ng/μl *Peft-3::cas9* (Addgene ID #46168 ^42^, 50-100 ng/μl u6::sgRNA with sequences targeted against either *cxTi10816, dhc-1, ebp-1, ebp-2* or *ebp-3,* 30-50 ng/μl; of the repair template, 50 ng/μl PAGE-purified *pha-1* repair oligonucleotide (Integrated DNA technologies), 60 ng/μl pJW1285 (Addgene ID #61252 ^75^), and 2.5 ng/μl *Pmyo-2::tdtomato* as a co-injection marker. Animals were grown for 3-5 days at either 20 or 25°C after injection, and transgenic progeny was selected based on either expression of tdTomato in the pharynx or survival at the non-permissive temperature (25°C). Subsequent assessment of genome editing events was carried out by either visual inspection using a wide-field fluorescence microscope and/or PCR amplification using primers targeting the inserted FP and a genomic region situated outside of the range of homology arms in case of *dhc-1,* or sequences flanking the predicted cut sites as well as an internal control in case of the Δ*ebp-1/2/3* knock-out mutants. The contexts of PCR-confirmed edited genomic loci were further inspected by sequencing (Macrogen Europe).

### Quantification of embryonic lethality and total brood size

In two separate experiments, N2, SV1598, SV1619 and SV1803, or N2, SV1868, SV1872, SV1877 and SV1882 single L4-stage hermaphrodites were placed on OP50 feeding plates and kept at 20 °C. Animals were transferred to a new plate every day.

On each plate, embryonic lethality was scored after 24 hours, and brood size 48 hours after removal of the parent. Experiments were executed in quadruplicate.

### Microscopy

For time-and stream-lapse imaging embryos were dissected from adult worms on coverslips in 0.8x egg salts buffer 94 mM NaCl, 32 mM KCl, 2.7 mM CaCl2, 2.7 mM MgCl2, 4 mM HEPES, pH 7.5^76^, and subsequently mounted on 4% agarose pads. Live-cell SDCLM imaging of one-cell embryos was performed on a Nikon Eclipse Ti with Perfect Focus System, Yokogawa CSU-X1-A1 spinning disc confocal head, Plan Apo VC 60x N.A. 1.40 oil and S Fluor 100x N.A. 0.5-1.3 (at 1.3, for photo-ablation) objectives, Photometrics Evolve 512 EMCCD camera, DV2 two-channel beam-splitter for simultaneous dual-color imaging, Cobolt Calypso 491 nm (100 mW), Cobolt Jive 561 nm (100 mW) and Teem Photonics 355 nm Q-switched pulsed laser controlled with the ILas system (Roper Scientific France/ PICT-IBiSA, Institut Curie, used for photo-ablation), ET-GFP (49002), ET-mCherry (49008) and ET-GFPmCherry (49022) filters, ASI motorized stage MS-2000-XYZ with Piezo Top Plate, and Sutter LB10-3 filter wheel. The microscope was controlled with MetaMorph 7.7 software and situated in a temperature-controlled room at 20°C. For regular single- and dual-channel imaging experiments, images were acquired in either stream-lapse mode with 100 ms exposure, or time-lapse mode with 500 ms exposure and 5 second intervals, unless stated otherwise. Laser power was kept constant within experiments. For spindle bisection assays, spindles were imaged after photo-ablation in stream-lapse mode with 500 ms exposure time.

Simultaneous dual-color TIRF imaging of embryos was performed on a Nikon Eclipse Ti with Perfect Focus System, Nikon Apo TIRF 100x N.A. 1.49 oil objective, Photometrics Evolve 512 EMCCD camera, Optosplit III beam-splitter for simultaneous dual-color imaging, 488 nm (150 mW) and Cobolt Jive 561 nm (100 mW) lasers, ET-GFP (49002), ET-mCherry (49008) and ET-GFPmCherry (49022) filters, ASI motorized stage MS-2000-XY System for Inversted Microscope Nikon Te/Ti 2000, and Sutter LB10-3 filter wheel. Acquisition was controlled with MetaMorph 7.7 software and the setup was situated in a temperature-controlled room at 20°C. For single- and dual-channel imaging experiments, images were acquired in stream-lapse mode with 100 ms exposure time.

Single-color TIRF imaging of embryos was performed on either above-mentioned TIRF setup, or on an identical TIRF setup in which lasers were controlled by the Ilas-2 system (Roper Scientific France / PICT-IBiSA, Institut Curie), and image acquisition was controlled with MetaMorph 7.8 software.

Live-cell wide-field time-lapse differential interference contrast (DIC) microscopy imaging of embryos was performed on a Zeiss Axioplan upright microscope, with a 100x N.A. 1.4 PlanApochroma objective, controlled by AxioVision Rel 4.7 software, at an acquisition rate of 1 image per 2 seconds with constant exposure time and light intensity. Embryos were followed from pronuclear meeting until completion of the first division. Images acquired by SDCLM and TIRF microscopy were prepared for publication in ImageJ by adjusting brightness and contrast, subtracting background and frame averaging as described in figure legends.

### RNA-mediated interference

For RNAi experiments^77^, either the gonads of young adults were injected with double-stranded RNA targeting RNA molecules of interest *(perm-1* and *perm-1* + *lin-5)* and grown for 20 hours at 15°C, or L4 animals were grown on RNAi feeding plates for 48 hours at 15°C prior to imaging sessions *(lin-5, ebp-1/3, ebp-2).*

### UV laser spindle midzone severing

Severing of the mitotic spindle was performed as described previously^4^. AZ224, TH65, SV1874, SV1879 and SV1900 embryos were cultured and imaged at 20°C, and subjected to spindle bisection at anaphase onset, as judged by GFP::TBB-2 and TBA-2::YFP tubulin localization. Peak centrosome velocities upon spindle severing were subsequently tracked automatically using image intensity thresholding and the FIJI TrackMate plugin.

### Data analysis

All intensity profile measurements on SDCLM and TIRF data were generated using ImageJ. Analysis of the timing of key mitotic events and position of the nucleocentrosomal complex and centrosomes from DIC movies of N2, SV1868, SV1872, SV1877 and SV1882 strains was performed by hand using ImageJ. Quantifications of cortical MT residence time were performed by hand using the FIJI plugin MtrackJ. Speeds of EBP-2::GFP and mCherry::DHC-1 comets during metaphase were calculated based on angles of tracks in kymograph data generated in ImageJ with the KymoResliceWide plugin. Numerical data processing was performed using Excel 2011 (Microsoft). Line and bar graphs were generated using Prism 6 (GraphPad software, inc.).

## Acknowledgements

We thank all of the members of the van den Heuvel, Akhmanova and Boxem groups for helpful discussions and general support. We acknowledge Wormbase and the Biology Imaging Center at the Faculty of Sciences, Department of Biology, Utrecht University, and especially Ilya Grigoriev and Eugene Katrukha for support regarding the technical aspects of microscopy and data analysis. Some strains were provided by the *Caenorhabditis* Genetics Center (CGC), which is funded by NIH Office of Research Infrastructure Programs (P40 OD010440). This work was supported by a European Research Council (ERC) Synergy grant 609822 to A.A., and is part of program CW711.011.01 (SvdH) financed by the Netherlands Organization for Scientific Research (NWO).

## Author contributions

AA, RS and SvdH designed the study, analyzed the data and wrote the paper. RS performed the experiments.

